# Reusable immobilised quaternary ammonium particles reduce microbial and resistome burdens without promoting resistance selection during wastewater post-treatment

**DOI:** 10.64898/2026.03.26.714185

**Authors:** Marta Redondo, Uli Klümper, Ana Pereira, Luis Melo, Thomas Berendonk, Alan Elena

## Abstract

Wastewater treatment plants act as convergence zones for antibiotic residues, antibiotic-resistant bacteria (ARB), and antimicrobial resistance genes (ARGs), yet conventional processes are not designed to mitigate resistance dissemination from their effluents. While chemical disinfectants are generally effective, soluble quaternary ammonium compounds (QACs) can generate subinhibitory exposure gradients that promote resistance selection and co-selection both during treatment and after release into receiving waters. Here, we evaluate a contact-restricted alternative: benzyldimethyldodecyl ammonium chloride (BDMDAC) immobilised onto hydroxyapatite microparticles as a reusable and retainable post-treatment polishing strategy. Across single-strain assays, treated wastewater exposure, and experimental community evolution, immobilised BDMDAC-functionalised particles (BDMDAC-FPs) achieved concentration-dependent antimicrobial activity without detectable biocide leaching. Optimal exposure (200 mg/L, 4 h) resulted in a ~5.5 log reduction in total bacterial abundance and removal of clinically relevant ARGs. Antimicrobial efficacy was retained after one reuse cycle, supporting operational stability. Plasmid-borne QAC ARGs did not confer protection, and no enrichment of qac-associated or non-QAC ARGs was observed. Conjugation assays demonstrated suppression of horizontal gene transfer even under suboptimal exposure, and mobility-associated markers remained stable or declined during long-term community incubation. Collectively, the data support a contact-restricted mechanism in which antimicrobial pressure is spatially confined to the particle interface, generating high local lethality while limiting diffuse subinhibitory exposure. This spatial confinement decouples antimicrobial efficacy from classical disinfectant-driven resistance selection and mobility amplification. Immobilised BDMDAC-FPs therefore provide a mechanistically distinct and evolution-conscious framework for wastewater polishing technologies.

## 1. Introduction

Wastewater treatment plants (WWTP) play a major role in water sanitation, especially in urban areas where wastewater can carry industrial, household, and nosocomial effluents. Their primary function is the removal of organic matter, nutrients, and chemical contaminants that pose threats to environmental, human, and animal health (e.g., personal care products, analgesics, and hormones)^1–3^. However, beyond these conventional targets, untreated wastewaters also carry antibiotics, alongside biological pollutants including pathogens, antibiotic-resistant bacteria (ARB), and antimicrobial resistance genes (ARGs), sometimes in very high concentrations^4,5^. As a result, wastewater treatment systems function not only as sanitation infrastructures but also as convergence zones where antibiotic residues, ARB, and mobile ARGs co-occur under conditions favourable to selection and genetic exchange^4^. WWTPs are not specifically designed to remove or mitigate these biological contaminants^6^, and even when properly functional, are not always able to fully eliminate these contaminants, leading to the release of pathogens, ARB, and ARGs into receiving water bodies^7–10^.

The presence of ARB and ARGs can have deleterious impacts on multiple levels: I) ARB can establish themselves in receiving water bodies, transforming these environments into reservoirs for AMR^11^; II) ARGs released into these systems can be taken up by naturally occurring bacteria, contributing to the emergence of new resistant populations^12,13^; and III) the co-occurrence of resistant and susceptible bacteria under selective pressures can facilitate horizontal gene transfer, promoting the accumulation and dissemination of multi-drug resistance trait^14^. Such contamination creates multiple environmental exposure pathways, including animal ingestion through drinking water^15^, direct human contact during recreational activities^16^, and indirect transmission via crop irrigation or potable water sources^17^, collectively contributing to the broader dissemination of wastewater-borne AMR. It is because of this that post-treatment strategies aimed at further decreasing biological contaminant loads are highly valuable.

Quaternary ammonium compounds (QACs) are among the most widely used disinfectants globally due to their broad-spectrum antimicrobial activity, chemical stability, and cost-effectiveness^18^. They are extensively applied across clinical, industrial, agricultural, and domestic settings, resulting in their continuous release into wastewater streams^19,20^. Their antimicrobial mode of action is primarily based on membrane disruption and leakage of intracellular contents, making them effective against a wide range of Gram-positive and Gram-negative bacteria^21^. Consequently, QACs are frequently detected in wastewater treatment systems and receiving environments, where their persistence contributes to ongoing antimicrobial selective pressures^22^.

Chemical disinfection remains an attractive post-treatment strategy due to its rapid antimicrobial action, operational simplicity, and scalability compared to more energy-intensive or infrastructure-demanding approaches such as advanced oxidation, membrane filtration, or ultraviolet irradiation^23^. Among chemical disinfectants, QACs are particularly appealing given their stability and broad-spectrum efficacy. However, when applied in soluble form, their environmental bioavailability and persistence can promote resistance selection and unintended ecological exposure^20,24^.

To overcome this, we previously developed and tested the potential of benzyldimethyldodecyl ammonium chloride (BDMDAC), a QAC representative, immobilised onto hydroxyapatite microparticles to act as a simple and time-effective post-treatment method for bacterial inactivation^9^. This immobilisation strategy successfully reduced bacterial concentrations under laboratory conditions while preventing detectable biocide leaching (LoD <15 ppm), supporting the production of chemically residue-free treated water^25^. Despite these promising initial insights, many questions regarding the potential of BDMDAC-functionalised particles (FPs) for ARB, ARG and pathogen mitigation remained open. In this work, we aimed to further characterise the performance and safety of BDMDAC-FPs as a wastewater post-treatment strategy. Specifically, we sought to: I) determine optimal particle exposure parameters for effective bacterial inactivation; II) evaluate antimicrobial efficacy across microorganisms displaying differing intrinsic and acquired resistance backgrounds; III) assess the functional reusability of particles as a determinant of operational feasibility; IV) investigate whether particle exposure promotes or suppresses horizontal gene transfer; and V) evaluate treatment performance in treated wastewater, including impacts on microbial community composition, pathogen-associated taxa, antimicrobial resistance gene abundance, and mobility determinants. Establishing these performance characteristics is essential to determine whether immobilised BDMDAC particles meet the functional and safety requirements necessary for integration into wastewater treatment infrastructures. Particular importance lies in achieving sustained antimicrobial efficacy without promoting pathogen enrichment, resistance selection, or enhanced mobility potential within treated effluents.

## 2. Methods

### 2.1 Synthesis of BDMDAC-functionalised particles

To generate immobilised disinfectant particles suitable for controlled antimicrobial testing, BDMDAC was immobilised onto hydroxyapatite microparticles using a layer-by-layer electrostatic assembly approach as described previously^25^. Briefly, hydroxyapatite microparticles (5.0 ± 1.0 µm; Fuidinova, Portugal) were sequentially coated with alternating charged polymers to enable stable BDMDAC attachment: Particles were first exposed to a 1 mg/mL polyethyleneimine (PEI) solution for 30 minutes, followed by suspension in a 1 mg/mL polystyrene sulfonate (PSS) solution for an additional 30 minutes. The resulting particles were subsequently incubated in a 1 mg/mL BDMDAC solution under stirring for 30 minutes to allow electrostatic immobilisation of the quaternary ammonium compound (QAC). Between each coating step, particles were recovered and washed twice to remove excess reagents. All coating and washing steps were performed in 0.1 M borate buffer (pH= 9). After functionalisation, particles were dried overnight at 80 °C and stored at 4 °C until use. For antimicrobial experiments, working suspensions were prepared by resuspending 0.220, 0.441, 0.882, and 1.764 g of dried functionalised particles in 100 mL of treatment solution, corresponding to final BDMDAC-equivalent concentrations of 25, 50, 100, and 200 mg/L, respectively.

### 2.2 Bacterial strains and growth inhibition assays

To determine the antimicrobial efficacy of BDMDAC-FPs and evaluate potential effects of intrinsic and plasmid-mediated QAC resistance, growth inhibition assays were performed using defined model organisms.

*Escherichia coli* MG1655 and *Pseudomonas putida* KT2442 were selected due to their environmental relevance, differing intrinsic tolerance to QACs, and suitability for conjugation assays. Isogenic derivatives carrying plasmid-borne QAC resistance determinants, plasmid RP4^26^ harbouring the functional efflux gene *qac*E and plasmid pKJK5^27^ carrying the truncated integron-associated variant *qacΔE*) were included to assess whether horizontally acquired disinfectant resistance alters susceptibility to immobilised BDMDAC. The *qacE* gene encodes a membrane-associated efflux pump conferring tolerance to soluble QACs, while *qacΔE* represents a widely distributed integron-associated derivative frequently co-localised with additional antimicrobial resistance determinants^28,29^. Inclusion of these plasmids, therefore, enabled evaluation of both functional disinfectant resistance and integron-associated resistance backgrounds under particle exposure.

Overnight cultures of the two strains with and without plasmids were grown in LB broth and adjusted to an OD_600_ corresponding to approximately 1 × 10^7^ CFU/mL. *E. coli* cultures were exposed to 25 and 100 mg/L BDMDAC-FPs, while *P. putida* cultures were exposed to 50 and 200 mg/L BDMDAC-FPs. Concentrations were selected based on prior characterisation of particle performance^25^, while untreated cultures served as controls.

All assays were conducted in agitation (100 rpm) at 37 °C. OD_600_ was recorded every 5 minutes over a 24-hour period using a microplate reader. Growth parameters, including lag phase duration, maximum growth rate, and carrying capacity, were calculated using the growthcurver package (v0.3.1) in R^30^. Statistical differences between treatment groups were assessed using Kruskal Wallis’ test followed by Dunn’s test.

### 2.3 Particle reuse assays

To evaluate the operational stability and reusability of BDMDAC-FPs as a determinant of economic and practical feasibility, particles were recovered and re-applied in sequential antimicrobial assays.

Following completion of growth inhibition experiments, BDMDAC-FPs were collected and washed with sterile tetrasodium pyrophosphate (TSPP) buffer to detach adhered biomass, followed by three washes with sterile saline solution. Washed particles were dried overnight at 37 °C, weighed to ensure similar exposure concentrations, and subsequently reused under the same exposure conditions described in Section 2.2. Up to two consecutive reuse cycles were performed. Antimicrobial efficacy after each reuse cycle was assessed using identical growth inhibition assays as described above. Statistical differences between treatment groups were assessed using Kruskal Walli’s test followed by Dunn’s test.

### 2.4 Conjugation assays

To determine whether BDMDAC-FP exposure influences plasmid-mediated horizontal gene transfer, conjugation assays were performed under suboptimal, partially inhibitory, and fully inhibitory particle exposure conditions.

For the *E. coli* system, a donor chromosomally-tagged with *mCherry* and kanamycin (KAN)-resistance carrying the conjugative, tetracycline (TET)-resistance plasmid pKJK5::*gfp*^31^ was paired with a gentamicin (GEN)-resistant, non-fluorescent recipient strain^32^. Overnight cultures were grown in LB broth supplemented with the appropriate antibiotics at 37 °C until reaching an OD_600_ of approximately 1. Cultures were diluted 1:10 in fresh LB, and donor and recipient were mixed at a 1:1 volume ratio in sterile glass containers. Four exposure conditions were tested: i) 25 mg/L BDMDAC-FPs (suboptimal), ii) 50 mg/L BDMDAC-FPs (partially inhibitory), iii) 100 and 200 mg/L BDMDAC-FPs (fully inhibitory for *E. coli* and *P. putida*), iv) no particle addition (control).

Mixtures were incubated for 4 hours at 37 °C with agitation (100 rpm). Following incubation, particles were allowed to sediment, and the supernatant was recovered. To evaluate potential surface-associated conjugation, particles were washed with TSPP buffer as described in Section 2.3, and the eluate was collected separately. Serial dilutions of supernatant and particle eluate were plated on LB agar supplemented with KAN (50 µg/mL), TET (10 µg/mL), and GEN (20 µg/mL) for transconjugant selection. Donor and recipient populations were quantified using selective single-antibiotic plates. The detection limit of the assay was (10 CFU/mL). All assays were performed in 5 biological replicates.

The same experimental design was applied to the *P. putida* system^31^ except that the recipient was rifampicin (RIF) resistant^33^. Selective plating for transconjugants was performed using LB agar supplemented with KAN (50 µg/mL), TET (10 µg/mL), and RIF (100 µg/mL).

Statistical differences between treatment groups were assessed using Student’s t-test.

### 2.5 Wastewater exposure experiments and quantitative PCR analysis

To evaluate the antimicrobial and resistance-mitigating performance of BDMDAC-FPs under environmentally relevant conditions, treated wastewater effluent was subjected to controlled particle exposure followed by molecular quantification of bacterial and resistance determinants.

#### 2.5.1 Wastewater exposure

Treated wastewater samples were collected in triplicate from an urban wastewater treatment plant (Dresden-Kaditz – December 2024). For each replicate, 100 mL of effluent was exposed to BDMDAC-FPs at final concentrations of 50 mg/L (partially inhibitory) and 200 mg/L (fully inhibitory) for 4 hours under agitation (150 rpm) at room temperature. Untreated wastewater samples processed in parallel served as controls. Following exposure, particles were separated by decantation. The supernatant of treated samples and untreated controls was filtered through 0.2 µm polycarbonate membrane filters (Sartorius) to collect cellular biomass. Filters were stored at −20°C until DNA extraction. Extracellular DNA present in the filtrate was concentrated using Macrosep centrifugal purification tubes with a 10 kDa molecular weight cut-off and subsequently purified using silica-based DNA extraction columns (Qiagen).

#### 2.5.2 DNA extraction and quantitative PCR

Total DNA from membrane filters was extracted using the DNeasy PowerSoil Pro kit (Qiagen – Hilden, Germany) according to the manufacturer’s instructions. DNA concentration and purity were assessed using fluorometric assays (Qubit – ThermoFisher Scientific, Massachusetts, USA).

Absolute bacterial abundance was quantified by targeting the 16S rRNA gene^34^. A six-point standard curve was generated using the pNORM plasmid as standard (http://www.norman-network.net/). Only amplification reactions with efficiencies between 90–110% and R² ≥ 0.99 were considered valid for quantification. All reactions were performed in 6 technical replicates across 5 biological replicates. Statistical differences in gene abundance between treatment groups were assessed using Tukey’s honest significant difference (HSD) on log transformed data. Primer sequences and amplification conditions are provided in Supplementary Table 1.

### 2.6 Community evolution

To evaluate the longer-term ecological impact of BDMDAC-FP exposure on wastewater microbial communities and associated resistance determinants, evolution experiments were conducted under repeated exposure conditions followed by 16S rRNA gene sequencing and high-throughput qPCR analysis.

#### 2.6.1 Community evolution setup

Treated wastewater (500 mL per replicate) was filtered through 0.2 µm membrane filters to collect microbial biomass. Filters were incubated overnight in synthetic wastewater medium (Composition provided in Supplementary Table 2) at room temperature to allow community recovery. Resulting cultures were homogenised and divided into 40 mL aliquots, which were exposed to 0 mg/L (control), 50 mg/L, or 200 mg/L BDMDAC-FPs. Cultures were incubated at room temperature under agitation (160 rpm) for 7 days. Fresh synthetic wastewater medium was supplied every 48 hours. All treatments were performed in quintuplicate.

At day 7, the complete volume of each culture was filtered through 0.22 µm polycarbonate membranes. Total DNA was extracted using the DNeasy PowerSoil Pro kit (Qiagen) according to the manufacturer’s instructions.

#### 2.6.2 16S rRNA gene amplification and sequencing

To determine community composition, the V3–V4 region of the 16S rRNA gene was amplified using previously validated primers^34^. Amplicon libraries were prepared using (library preparation kit placeholder) and sequenced using 150 bp paired-end chemistry on an Illumina NovaSeq 6000 platform. Raw reads were quality-checked using FASTQC^35^ and trimmed using BBDuk^36^ to remove adapters and retain reads with Q > 30. High-quality reads were processed using a DADA2-based workflow in R^37^ to generate amplicon sequence variants (ASVs). Taxonomic assignments were performed against the SILVA reference database (version 138.2)^38^. Community dissimilarities were calculated using Bray–Curtis distances and visualised using non-metric multidimensional scaling (NMDS). Statistical differences between treatment groups were assessed using the analysis of similarities (ANOSIM) test on a Bray-Curtis distance matrix. Raw sequencing reads have been deposited in the NCBI Sequence Read Archive (SRA) under BioProject accession number PRJNA1431717.

#### 2.6.3 Pathogen-associated genus analysis

To assess whether BDMDAC-FP exposure resulted in enrichment of potentially pathogenic taxa, genus-level taxonomic profiles were screened against a curated list of bacterial genera containing recognised human pathogens^39^, following previously established methodology^39^. For each sample, the cumulative relative abundance of reads assigned to pathogen-associated genera was calculated as the proportion of total classified reads. Statistical differences between treatment groups were assessed using Kruskal Wallis’ Test followed by Dunn’s test.

#### 2.6.4 High-throughput quantitative PCR analysis

To evaluate the impact of prolonged BDMDAC-FP exposure on AMR and mobility determinants, chip-based high-throughput qPCR was performed targeting 32 genes, including major antimicrobial resistance genes, QAC resistance determinants, and plasmid replicon markers (Primer sequences provided in Supplementary Table 3). Extracted DNA from each evolution replicate was used as template for parallelised amplification reactions. Only amplification curves exhibiting a single melt peak, efficiencies between 1.8 and 2.2, and Ct values below 27 were included in downstream analyses. Relative gene abundance was calculated by normalising target gene Ct values to 16S rRNA gene Ct values using the ΔCt method^40^.

Statistical differences between treatments were evaluated using Kruskal Wallis’ test followed by Dunn’s test, all values were corrected for multiple comparisons using Bonferroni’s correction.

## 3. Results

### 3.1 Immobilised quaternary ammonium particles exert broad antimicrobial activity independent of resistance background

To evaluate the feasibility of immobilised quaternary ammonium particles as a wastewater post-treatment strategy, we first assessed their antimicrobial performance under controlled exposure conditions. Particular attention was given to organisms displaying distinct intrinsic disinfectant tolerance and to the potential influence of plasmid-borne quaternary ammonium resistance determinants.

Particle exposure resulted in pronounced, concentration-dependent growth inhibition across all tested strains. Importantly, the presence of plasmid-encoded QAC resistance genes did not confer measurable survival or growth advantages under any exposure condition.

In detail, exposure of *Escherichia coli* to 25 mg/L BDMDAC-FPs significantly impaired growth kinetics and carrying capacity relative to untreated controls (Figure 1A; Supplementary Table 4) (carrying capacity *E. coli* _25mg/mL_ 1.59 ± 0.05, *E. coli* _Control_ 3.44 ± 0.05, Dunn’s test, *p*<0.0001). However, residual growth was still observed after 20 hours of exposure, indicating suboptimal antimicrobial activity at this concentration over longer time periods. Increasing particle exposure to 100 mg/L resulted in complete growth suppression, with no detectable growth phase throughout the experimental period (Figure 1a; Supplementary Table 4) (carrying capacity *E. coli* _100mg/mL_ 0.29 ± 0.05 Dunn’s test, *p*<0.0001).

**Figure 1:**
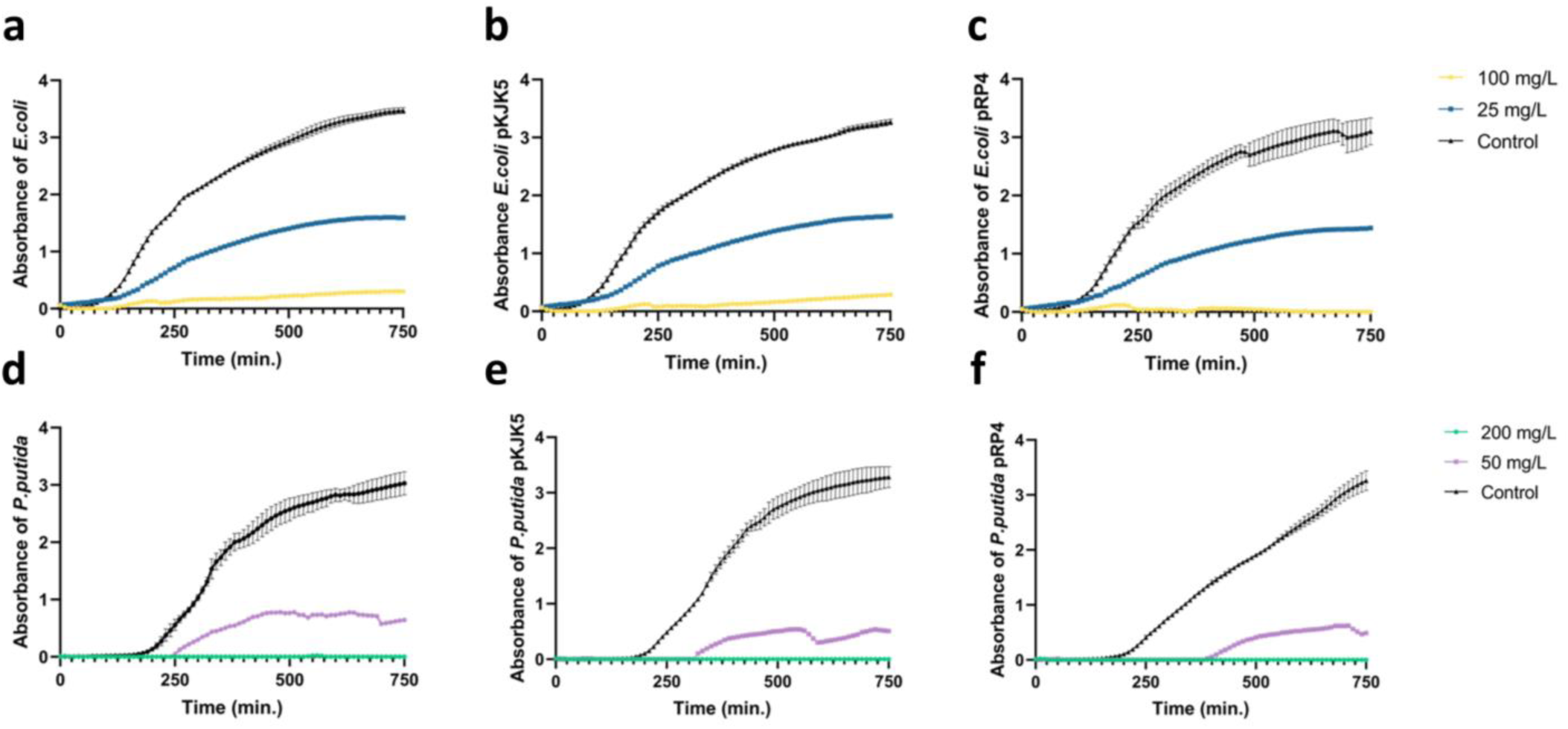
Photometric measurements of growth curves of different species and strains exposed to different concentrations of BDMDAC-FP. a) *E. coli* MG1655, b) *E. coli* MG1655 (pKJK5 - *qacΔE*), c) *E. coli* MG1655 (pRP4-*qacE*), d) *P. putida* KT2442, e) *P. putida* KT2442 (pKJK5 - *qacΔE*) and f) *P. putida* KT2442 (pRP4 - *qacE*). Bars represent standard deviation.

To assess whether horizontally acquired QAC resistance could mitigate particle activity, we evaluated isogenic *E. coli* strains harbouring plasmids encoding quaternary ammonium resistance determinants. Two broad-host-range model plasmids were selected: RP4 harbouring the functional efflux gene *qacE*, and pKJK5 carrying the truncated integron-associated variant *qacΔE*. The *qacE* gene encodes a membrane-associated efflux pump conferring tolerance to soluble quaternary ammonium compounds^28^, while *qacΔE* represents a widely distributed derivative commonly embedded within class 1 integron structures and frequently co-localised with additional AMR determinants^29^. The use of these plasmids therefore enabled assessment of both functional disinfectant resistance and integron-associated resistance backgrounds under particle exposure. Despite their carriage, growth inhibition patterns remained indistinguishable from those observed in plasmid-free strains across all tested concentrations (Figure 1b–c; Supplementary Table 4). Neither plasmid conferred measurable protection against immobilised BDMDAC, indicating that resistance determinants typically associated with soluble QAC exposure do not mitigate particle-mediated antimicrobial activity.

In contrast, *Pseudomonas putida* displayed higher intrinsic tolerance, requiring a minimum exposure of 50 mg/L to observe measurable growth inhibition (Figure 1d). At this concentration, reduced growth rates and carrying capacities were detected relative to controls (Supplementary Table 4) (carrying capacity *P. putida* _Control_ 2.93 ± 0.17, *P. putida* _50mg/L_ 0.57 ± 0.05, Dunn’s test, *p*<0.0001). Complete growth suppression was achieved at 200 mg/L, where no proliferation was observed over the experimental timeframe (Figure 1d; Supplementary Table 4) (carrying capacity *P. putida* _200mg/mL_ 0.00 ± 0.00, Dunn’s test, *p*<0.0001). As with *E. coli*, the presence of either of the QAC resistance plasmids did not modify susceptibility patterns (Figure 1e–f), confirming that plasmid-encoded resistance did not confer an advantage even in intrinsically resilient hosts.

These results highlight species-specific tolerance thresholds while confirming robust antimicrobial efficacy under operationally realistic exposure conditions. Based on cross-species performance, 200 mg/L BDMDAC-FPs with a minimum exposure time of four hours was selected as a generalised full-inhibition treatment parameter for subsequent risk mitigation experiments, with 50 mg/L serving as an incomplete inhibition treatment.

Having established antimicrobial efficacy under defined operational conditions, we next assessed whether functionalised particles retained activity following reuse, a critical determinant of technological feasibility and cost efficiency for large-scale wastewater post-treatment deployment.

### 3.2 Functionalised particles retain antimicrobial efficacy following reuse

To evaluate the operational feasibility and cost efficiency of BDMDAC-FPs, we next assessed whether functionalised particles retained antimicrobial activity following reuse. Recycled BDMDAC-FPs were thus recovered from prior antimicrobial assays, washed to remove adhered biomass, and reintroduced into bacterial growth inhibition experiments under previously defined suboptimal and optimal exposure conditions.

Particle reuse following a single operational cycle resulted in antimicrobial performance comparable to that observed with freshly prepared particles for both strains (Figure 2a; Supplementary Table 5). Suboptimal exposure concentrations continued to induce delayed growth and reduced carrying capacities relative to untreated controls (*E. coli* _25mg/mL_ = 1.30 ± 0.05, *P. putida* _50mg/mL_ = 0.00), while concentrations previously defined as optimal maintained rapid and complete growth suppression (carrying capacity *E. coli* _Control_ = 3.23 ± 0.04, *E. coli* _100mg/mL_ = 0.05 ± 0.001, *p*<0.0001 Dunn’s test. carrying capacity *P. putida* _Control_ = 3.15 ± 0.02, *P. putida* _200mg/mL_ = 0.08 ± 0.004, *p*<0.0001 Dunn’s test.). As observed with unused particles, plasmid carriage did not confer measurable survival advantages under any reuse condition.

**Figure 2:**
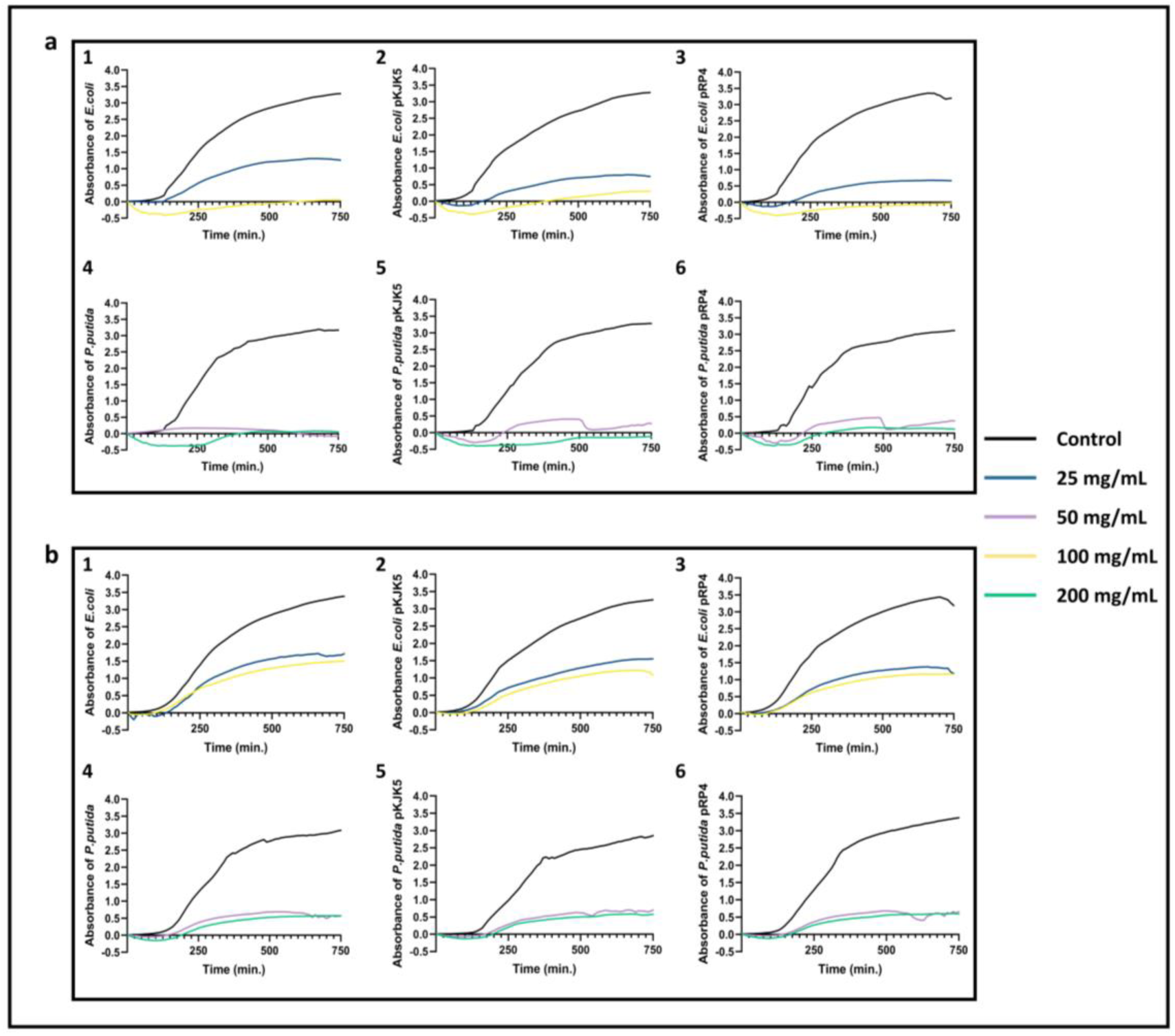
Photometric measurements of growth curves of different strains exposed to BDMDAC-FP reused one (a) and two (b) times. 1) *E. coli* MG1655, 2) *E. coli* MG1655 (pKJK5 - *qacΔE*), 3) *E. coli* MG1655 (pRP4-*qacE*), 4) *P. putida* KT2442, 5) *P. putida* KT2442 (pKJK5 - *qacΔE*) and 6) *P. putida* KT2442 (pRP4 - *qacE*).

A second reuse cycle, however, revealed slightly reduced efficacy was detected for both tested species. Under these conditions, survival was observed even at exposure concentrations previously defined as fully inhibitory, but resulting bacterial numbers after 12.5 hours still remained significantly reduced (*E. coli* _25mg/mL_= 59.76 ± 0.05% reduction, *E. coli* _100mg/mL_= 98.54 ± 0.05%, *P. putida* _50mg/mL_=100 ± 0.19% and *P. putida* _200 mg/mL_=97.62 ± 0.17% - Figure 2b. All *p*<0.001, Tukey’s HSD). Still, this minor partial loss of activity indicates that repeated reuse can compromise particle functionality and therefore represents an operational boundary for sustained antimicrobial performance, particularly when targeting disinfectant-tolerant environmental taxa.

Together, these findings support the feasibility of at least one reuse cycle without compromising antimicrobial efficacy, while in subsequent reuse cycles effective concentrations might need to be increased to maintain efficacy. This reuse potential improves the economic and operational viability of immobilised BDMDAC-FPs as a wastewater post-treatment strategy.

### 3.3 Particle exposure suppresses horizontal gene transfer despite bacterial aggregation and stress

Having established antimicrobial efficacy and operational feasibility, we next assessed whether particle deployment could influence AMR dissemination processes. Previous observations suggest that Immobilised BDMDAC particles attract bacteria and extracellular DNA to their surface through ionic interactions^25^. In combination with sublethal antimicrobial exposure, such aggregation and stress conditions could theoretically enhance plasmid exchange and promote horizontal gene transfer^41^, thereby selecting for a more mobile and higher-risk resistome^42^. To evaluate this possibility, we conducted conjugation assays under suboptimal particle exposure conditions that allowed donor and recipient survival while maintaining particle contact. Across all tested exposure conditions, particle treatment did not enhance plasmid transfer. Instead, horizontal gene transfer was consistently suppressed, even under suboptimal antimicrobial exposure. For the *E. coli* system, a gentamicin-resistant *E. coli* MG1655 strain was used as the recipient, while an *E. coli* plasmid donor carrying the conjugative plasmid pKJK5::*gfp,* conferring tetracycline and kanamycin resistance, was employed. A concentration of 25 mg/L BDMDAC-FPs was selected as suboptimal exposure, while 50 mg/L and 200 mg/L represented partial- and full-inhibitory concentrations, respectively. Donor and recipient cultures were mixed in equal proportions and allowed to conjugate for 4 hours in LB with or without BDMDAC-FPs exposure.

Untreated *E. coli* controls yielded 1.46×10^2^ ± 0.22 CFU/mL transconjugants (Figure 3a), confirmed by selective plating and fluorescence microscopy, while donor and recipient numbers stayed consistent from 0 to 4 hours (Donor: 4.60 ×10^4^ ± 0.65 CFU/mL vs. 4.25 ×10^4^ ± 0.21 CFU/mL, Recipient: 4.33 ×10^4^ ± 0.44 CFU/mL vs. 4.01 ×10^4^ ± 0.14 CFU/mL; Student’s t-test, both *p*>0.05). Suboptimal particle exposure resulted in substantial and significant reductions in survival of both donor (51.86±7.08%, Student’s t-test *p*<0.01) and recipient (49.35 ± 4.87%, Student’s t-test *p*<0.01) after 4 hours. Despite the survival of both mating partners, transconjugant numbers remained consistently below the detection limit of 10 CFU/mL following particle exposure (Figure 3a). Exposure to partially inhibiting particle concentrations (50 mg/L) also significantly reduced recipient survival while eliminating detectable donor populations, again yielding no detectable transfer events (Figure 3a). Fully inhibitory concentrations (200 mg/L) resulted in the complete absence of donor, recipient, and transconjugant cells.

**Figure 3:**
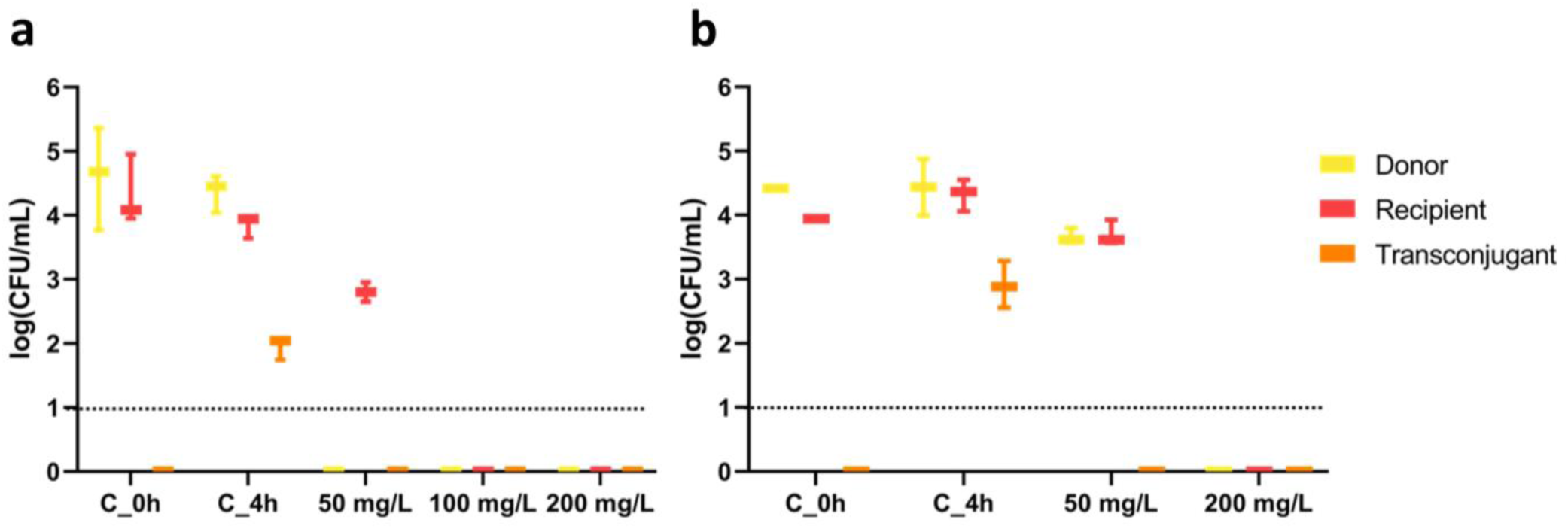
Conjugation assay results for a) *E. coli* and b) *P. putida*. Reported values belong to supernatants. No growth was obtained in particle eluate. Dotted lines represent the limit of detection of 10 CFUs/mL.

Comparable results were obtained in the *P. putida* conjugation system. Here, a rifampicin-resistant recipient strain was paired with a pKJK5::*gfp* donor, with untreated controls yielding approximately 10³ CFU/mL confirmed transconjugants (Figure 3b). While under partially-inhibiting particle exposure (50 mg/L), both donor and recipient populations remained culturable at 3.66×10⁴ ± 0.10 and 3.70×10^4^ ± 0.16 CFU/mL, still no transconjugants were detected (Figure 3b). As observed in the *E. coli* system, optimal particle exposure (200 mg/L) eliminated detectable donor, recipient, and transconjugant populations.

Throughout, no growth was obtained from particle eluates in any experimental condition, indicating that surface-associated conjugation did not occur despite bacterial aggregation at the particle interface.

We thus demonstrated that particle exposure does not promote plasmid-mediated resistance dissemination at the single-strain level.

### 3.4 Particle exposure reduces microbial load in treated wastewater

Having established antimicrobial efficacy, operational feasibility, and suppression of plasmid-mediated transfer in controlled systems, we next evaluated particle performance in a real-world wastewater context. Treated effluent collected from an urban wastewater treatment plant was exposed to BDMDAC-FPs under previously defined partly and fully inhibitory exposure conditions. To assess bulk antimicrobial performance in this complex microbial matrix, total bacterial abundance was quantified via qPCR targeting the 16S rRNA gene.

Particle exposure resulted in a pronounced reduction in microbial load relative to untreated controls. Fully inhibitory treatment conditions for the single strains (200 mg/L BDMDAC-FPs) achieved a significant 5.67 ± 0.393 log reduction in 16S rRNA gene copies in the complex community (Figure 4a; Tukey’s HSD, *p*<0.001), while partial inhibitory exposure (50 mg/L BDMDAC-FPs) still resulted in a 5.208 ± 0.226 log decrease in bacterial abundance (Figure 4a, Tukey’s HSD, *p*<0.001).

**Figure 4:**
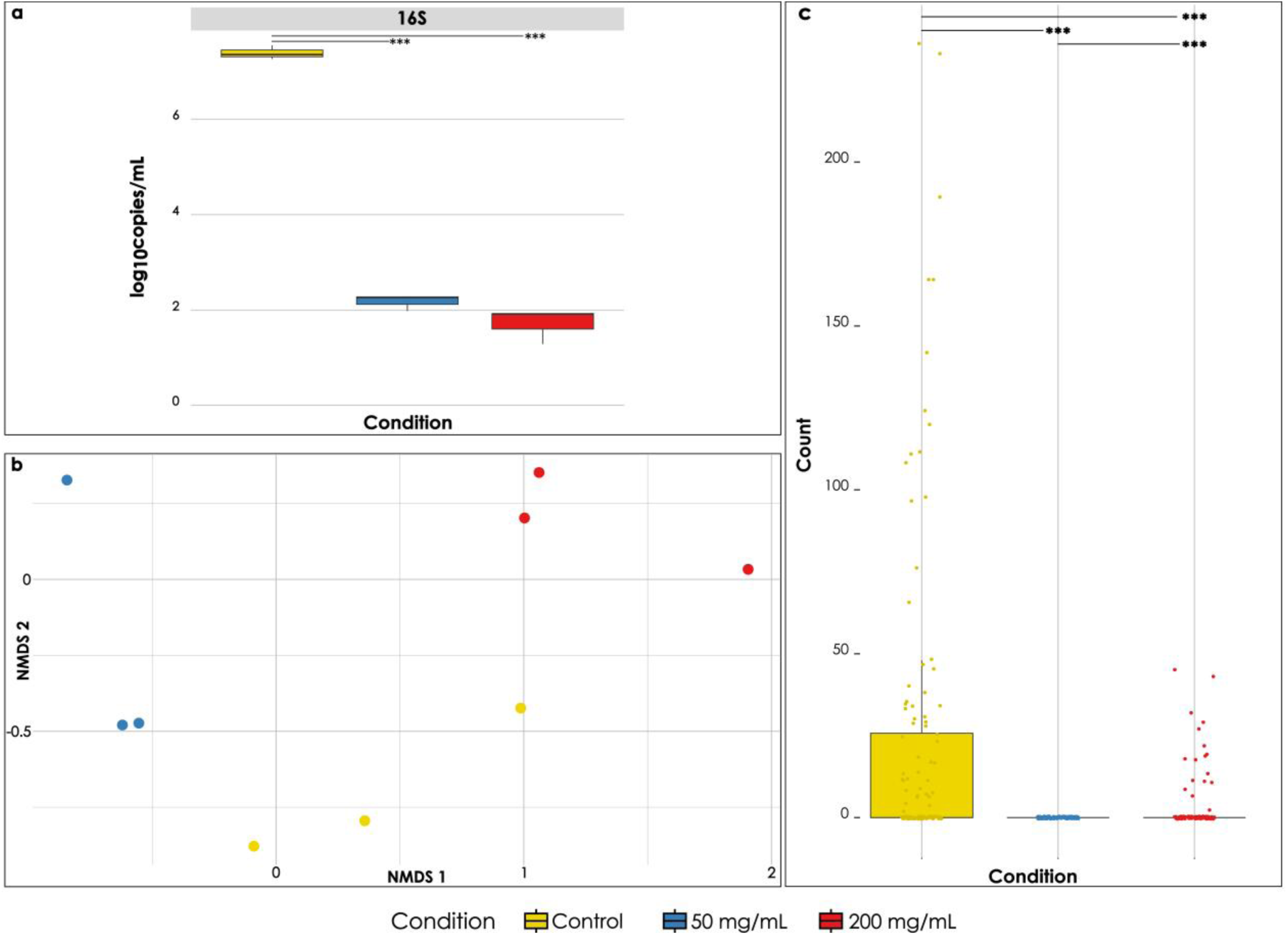
Overall bacterial removal (a), community divergence induced by long-term BDMDAC exposure (b) and effect of BDMDAC on the selection of known pathogens (c). *** represent significant differences between treatments (*p*<0.001).

These findings confirm that BDMDAC-FPs retain antimicrobial activity in treated wastewater and effectively reduce overall microbial burden under environmentally relevant conditions.

### 3.5 Particle exposure restructures wastewater communities without pathogen enrichment

Given the pronounced antimicrobial activity observed in treated wastewater, we next evaluated whether particle exposure induced selective restructuring of microbial communities and whether such shifts carried implications for pathogen enrichment. Surface-associated growth and sublethal stress conditions are known to favour opportunistic and host-adapted taxa capable of tolerating immune-like pressures^43,44^. Enrichment of such organisms during treatment would represent a downstream human health risk, particularly if released into receiving environments^45^.

Community composition analysis based on 16S rRNA gene sequencing revealed clear structural divergence between treated and untreated systems. Ordination analysis using non-metric multidimensional scaling (NMDS) demonstrated that evolved communities exposed to BDMDAC-FPs clustered distinctly from both the source wastewater community and untreated incubation controls (Figure 4b; all *p* < 0.05, ANOSIM).

To assess whether these structural shifts translated into altered pathogenicity potential, genus-level community profiles extracted from 16S rRNA gene analysis were compared against a curated list of bacterial genera containing known human pathogens, following previously established methodology^39^. Relative sequence abundance assigned to pathogen-associated genera was quantified for each treatment condition. This revealed a reduction in the proportional representation of potentially pathogenic genera in both particle-treated communities (Figure 4c) compared to the untreated control (Control vs 50 mg/mL *p* _Bonferroni_<0.001; Control vs 200 mg/mL *p* _Bonferroni_=0.002). For example, the relative abundance *Pseudomonas*, a genus including ample opportunistic pathogens^46^, was reduced in particle-treated communities compared to untreated controls (58.21% and 96% reduction in treatment with 50 mg/mL and 200 mg/mL BDMDAC). In contrast, indicator analysis identified genera such as *Elizabethkingia*, *Mitsuaria*, *Vogesella*, and *Delftia* as characteristic of particle-exposed communities, none of which are recognised as major human or animal pathogens (Indicator value = 0.994, *p* _Bonferroni_=0.02; Indicator value= 1, *p* _Bonferroni_=0.0183; Indicator value = 1, *p* _Bonferroni_=0.00183; Indicator value = 0.985, *p* _Bonferroni_=0.0183,). Together, these shifts indicate that community restructuring under particle exposure does not favour pathogen enrichment.

Importantly, reductions in relative pathogen-associated abundance occurred alongside the previously observed ~5.5 log decrease in absolute bacterial load. This dual reduction indicates that particle exposure not only lowers the proportional representation of potentially pathogenic taxa but also substantially reduces their absolute abundance.

Having established that particle exposure does not enrich pathogenic taxa despite pronounced community restructuring, we next assessed whether resistance and mobility determinants were similarly affected at the resistome level.

### 3.6 Particle exposure reduces resistome abundance without evidence of co-selection

Having established that particle exposure does not enrich pathogenic taxa, we next assessed whether AMR and AMR-mobility determinants were subject to direct selection or co-selection under treatment conditions. Quaternary ammonium compounds are known to select for disinfectant resistance genes and may co-select antibiotic resistance determinants through shared mobile genetic elements^47^. Moreover, while single-strain conjugation assays demonstrated that particle exposure suppresses plasmid-mediated transfer, it remained necessary to evaluate whether these mobility-limiting effects translated to complex wastewater resistomes.

We first evaluated markers associated with direct disinfectant resistance selection. Relative abundance analysis of qac-ARGs revealed no enrichment following particle exposure. Instead, QAC resistance-associated determinants remained stable or declined in relative abundance across treatment conditions (Figure 5b), indicating that immobilised BDMDAC exposure does not select for disinfectant resistance within wastewater communities. For example, *qacA*, a gene commonly associated to benzalkonium chloride, showed a significant decrease in the communities treated with 50 mg/mL and 200 mg/mL BDMDAC-FPs (Dunn’s test, *p* _Bonferroni_<0.05).

**Figure 5:**
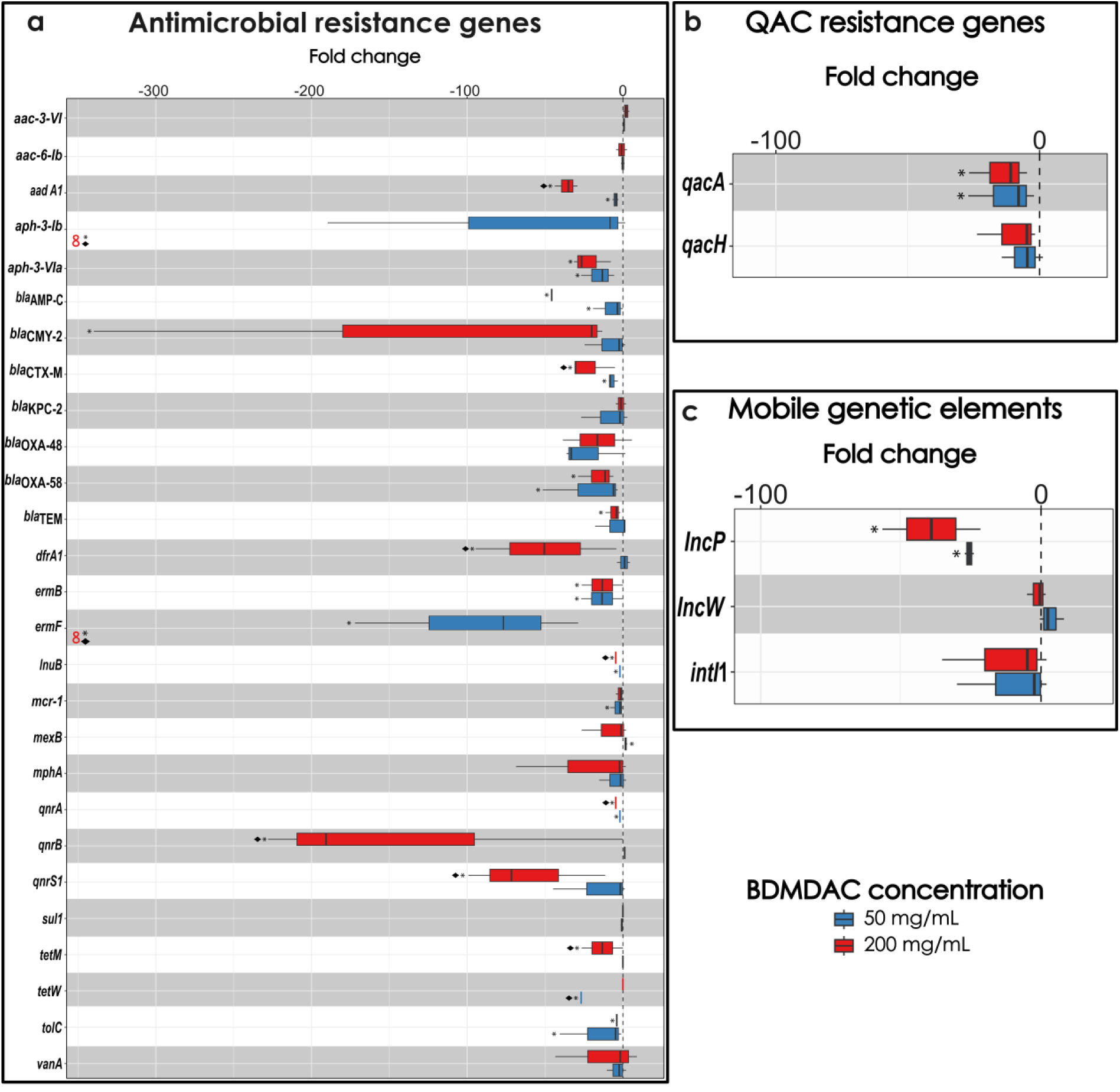
Fold change of relative abundance compared to the non-exposure control of ARGs (a), QAC resistance genes (b) and mobile genetic elements and related markers (c). Differences in means were assessed using the Kruskal-Wallis test followed by Dunn’s test. *p*-values were corrected for multiple comparisons using Bonferroni’s correction. * Represent significant differences between treatment and control (*p*<0,05) while ♦ represent significant differences between treatments.

We next assessed whether ARGs not conferring resistance to QACs were subject to co-selection. High-throughput qPCR profiling of a broad ARG panel for diverse antibiotic classes demonstrated widespread significant reductions in relative ARG abundance following treatment for 20 out of 27 tested ARGs, including clinically significant examples such as *mcr-1*, *bla*_CTX-M_ and *aph-(3’)-VI* (5, 22 and 21.9-fold; Tukey’s HSD, *p*<0.05), while no ARGs showed significant enrichment relative to untreated controls (Figure 5a). In several cases, for example *aadA1* and *qnrA*, ARG relative abundance declined progressively with increasing particle exposure, suggesting resistance gene deselection rather than co-selection under treatment pressure (Figure 5a; Tukey’s HSD, *p<0.05*).

To determine whether mobility potential was similarly affected, we quantified markers associated with horizontal gene transfer and genetic mobility. Integron-associated genes and plasmid-related determinants displayed stable (*incW*: −0.8 to 3.8-fold change – *intI*1: −34.87 to 2.4 fold change; both Tukey’s HSD, *p* _Bonferroni_>0.05) or significantly reduced relative abundance (*incP*; −25.2 to −38.7 fold change; Tukey’s HSD, *p* _Bonferroni_<0.05) across treatment conditions (Figure 5c). No mobility markers showed evidence of enrichment, indicating that particle exposure does not promote genetic exchange capacity at the community level. These findings align with the previously observed suppression of plasmid transfer in single-strain conjugation assays, collectively supporting the conclusion that particle exposure does not enhance AMR dissemination risk.

Finally, resistome reductions in relative abundance occurred in the context of the previously observed ~5.5 log decrease in total bacterial load. This concurrent decline in both microbial biomass and resistance gene representation indicates that particle exposure reduces the overall resistome burden within treated wastewater systems.

Collectively, these findings demonstrate that BDMDAC-FP exposure achieves effective microbial and resistome reduction while remaining functionally reusable and without promoting pathogen enrichment, resistance co-selection, or enhanced mobility potential across experimental scales.

## 4. Discussion

The present study demonstrates that immobilised BDMDAC-FPs achieve robust antimicrobial efficacy in wastewater systems while avoiding key risk pathways traditionally associated with soluble chemical disinfectants. Across controlled single-strain assays, complex wastewater exposure, and experimental community evolution, particle treatment consistently reduced microbial abundance, antimicrobial resistance gene load, and mobility-associated determinants without evidence of pathogen enrichment or resistance co-selection. Importantly, these effects were achieved in the absence of detectable biocide leaching, indicating that antimicrobial activity was spatially confined to the particle interface rather than distributed throughout the aqueous phase.

Chemical disinfection remains one of the most efficient strategies for microbial control^18^. However, the environmental deployment of soluble biocides, including QACs, has raised concerns regarding subinhibitory exposure, resistance selection, and unintended ecological impacts^18,24,47^. In soluble form, QACs are bioavailable throughout the water column, potentially generating diffuse selection gradients that favour tolerant organisms and co-select for AMR located on shared MGEs^47^. By contrast, immobilisation onto hydroxyapatite microparticles constrains BDMDAC bioavailability to direct contact events. This spatial restriction likely produces high local antimicrobial pressure at the particle surface while limiting persistent low-level exposure in the surrounding aqueous phase. The absence of enrichment of qac-associated ARGs and the widespread reduction of non-QAC resistance determinants observed here are consistent with a contact-restricted mechanism that reduces opportunities for classical disinfectant-driven selection and co-selection. This framework also provides a mechanistic explanation for the pronounced concentration dependence observed across experiments. Because antimicrobial activity is confined to the particle surface, sufficient particle density is required to ensure adequate contact frequency and surface area coverage for effective inactivation of the entire bacterial population. At lower particle concentrations, incomplete contact events likely permit survival of a fraction of cells, whereas higher concentrations increase the probability of lethal surface interactions across the community.

Species-specific differences in susceptibility further support this interpretation. *P. putida*, which is recognised for its surface adaptability and stress tolerance^48^, required higher particle concentrations to achieve complete growth suppression compared to *Escherichia coli*. Rather than reflecting reduced intrinsic sensitivity to BDMDAC, this pattern is consistent with differences in cell surface properties or aggregation behaviour that may influence particle–cell encounter dynamics. Under a contact-restricted antimicrobial model, organisms with enhanced structural resilience or altered surface interactions may require higher particle densities to ensure sufficient lethal interactions without increasing the amount of QAC immobilised per particle.

This contact-limited behaviour distinguishes immobilised BDMDAC from diffusible disinfectants, where antimicrobial activity scales with bulk-phase concentration rather than particle–cell encounter frequency^49^. By spatially confining antimicrobial pressure to the particle interface, immobilisation likely avoids the generation of diffuse subinhibitory exposure gradients that are frequently implicated in resistance enrichment and co-selection^47,50,51^.

The lack of protection conferred by plasmid-borne QAC resistance genes further supports this interpretation. Efflux-mediated tolerance mechanisms are effective against diffusible compounds but are unlikely to counteract high-intensity surface-confined interactions. This mechanistic distinction provides a plausible explanation for why integron-associated *qac* variants, frequently linked to multidrug resistance plasmids^47^, were not enriched under particle exposure. Consequently, immobilised BDMDAC did not trigger the selective dynamics commonly reported for soluble QAC exposure^47,50,51^.

Notably, particle treatment did not promote horizontal gene transfer, despite theoretical expectations that surface-associated aggregation and sublethal stress could enhance conjugation. Instead, plasmid transfer was consistently suppressed under both suboptimal and inhibitory exposure conditions. At the community level, mobility-associated markers, including integron and plasmid indicators, remained stable or declined during long-term exposure. These findings suggest that immobilised BDMDAC treatment does not amplify the dissemination potential of ARGs and may, in fact, reduce opportunities for genetic exchange by rapidly decreasing viable donor and recipient populations while maintaining high local antimicrobial pressure at the particle interface^52,53^.

In addition to suppressing conjugative transfer, particle exposure was not associated with detectable accumulation of extracellular DNA in treated wastewater. While photometric growth curves did not display a classical lysis-associated death phase, regrowth assays confirmed complete loss of viability under optimal exposure conditions, indicating bactericidal rather than merely bacteriostatic activity. Together, these observations are consistent with a rapid inactivation mechanism that does not involve extensive cell lysis and large-scale release of intracellular contents into the bulk phase. From an AMR risk perspective, this distinction is relevant, as lytic disinfection processes can increase the availability of extracellular resistance genes for potential uptake via natural transformation^54^. Although direct quantification of membrane disruption dynamics was beyond the scope of this study, the absence of detectable free DNA accumulation suggests that immobilised BDMDAC does not exacerbate secondary gene dissemination through extracellular DNA release.

Despite the pronounced ~5.5 log reduction in total bacterial abundance, particle exposure induced measurable restructuring of wastewater microbial communities during long-term incubation. Such compositional shifts are expected under strong antimicrobial pressure and reflect differential survival capacities across taxa^55^. Importantly, however, restructuring did not translate into enrichment of pathogen-associated genera. Instead, the overall relative abundance of genera containing opportunistic pathogens, declined under particle treatment, and no clinically dominant taxa emerged as treatment-associated indicators. This observation is critical, as selective enrichment of stress-tolerant pathogens has been reported under certain disinfection regimes and represents a key downstream human health concern^56^. Together, these patterns indicate that BDMDAC-FP exposure reduces both the proportional representation and the total load of potentially pathogenic organisms within treated wastewater. From a risk perspective, this dual effect is particularly relevant: while relative abundance informs the likelihood of post-treatment community regrowth^57^, absolute abundance influences propagule pressure into receiving environments and subsequent exposure pathways^58,59^. The absence of pathogen enrichment under strong antimicrobial selection therefore supports the safety profile of immobilised BDMDAC treatment in complex microbial systems.

At the resistome level, immobilised BDMDAC exposure did not induce enrichment of disinfectant resistance genes or evidence of ARG co-selection. Relative abundance analyses revealed stable or declining levels of QAC-associated determinants following treatment, reinforcing the absence of direct disinfectant selection observed in single-strain assays. This finding is particularly relevant given the frequent localisation of *qac* genes within class 1 integrons and multidrug resistance plasmids^47^, where soluble QAC exposure has been reported to drive co-selection dynamics^47^. Beyond QAC-associated markers, high-throughput profiling demonstrated widespread reductions in diverse ARGs across multiple classes, with no tested ARG exhibiting significant enrichment relative to untreated controls. In several cases, ARG abundance declined progressively with increasing particle concentration, consistent with resistance deselection under strong antimicrobial pressure rather than co-selection. Importantly, markers associated with genetic mobility, including integron and plasmid indicators, remained stable or decreased during long-term exposure. This community-level stability aligns with the experimentally observed suppression of plasmid conjugation in defined donor–recipient systems, indicating that immobilised BDMDAC does not enhance horizontal dissemination potential across biological scales.

Crucially, reductions in relative ARG abundance occurred in the context of a substantial absolute decrease in total bacterial biomass. This concurrent decline in both microbial load and resistance gene representation suggests that particle treatment reduces overall resistome burden rather than reshaping it toward a more mobile or high-risk configuration. Collectively, these findings contrast with selection patterns reported for diffusible disinfectants^24,51^ and support the concept that spatially confined antimicrobial exposure can achieve effective microbial control without amplifying resistance propagation pathways.

From an engineering perspective, the performance of BDMDAC-FPs compares favourably with established post-treatment approaches. Reported ARG removal efficiencies were within or above the range described for membrane bioreactors and conventional chlorination systems^60,61^, while avoiding continuous chemical dosing into the bulk phase. In contrast to oxidising disinfectants, which may release intracellular DNA and generate transformation-relevant substrates during lytic inactivation processes^62^, immobilised BDMDAC operates via spatially confined surface interactions. Similarly, while membrane-based systems rely on physical retention and can require substantial energy input and fouling management, particle-based treatment represents a modular contact strategy that could be implemented as a polishing step without major infrastructural modification. Importantly, the absence of detectable biocide leaching distinguishes this approach from soluble QAC application and reduces the likelihood of sustained environmental exposure.

Operationally, antimicrobial efficacy was retained after one reuse cycle, supporting the functional stability of the immobilised disinfectant layer. The partial decline in activity after repeated reuse is unlikely to result from chemical depletion, given the previously demonstrated absence of measurable BDMDAC release, and is more plausibly linked to cumulative surface masking or adsorption phenomena. Optimisation of particle regeneration strategies may therefore further enhance operational longevity and environmental benefit. Future work should investigate particle–cell interaction dynamics, membrane integrity during inactivation, and extracellular DNA release kinetics to more precisely define the molecular mechanism underlying contact-mediated killing.

While the present study provides multi-scale evidence across controlled and community-level systems, experiments were conducted under batch conditions and using a single wastewater source. Evaluation under continuous-flow operation and across diverse wastewater matrices will be required to fully assess robustness and scalability.

Beyond performance metrics, the relevance of effective ARG mitigation in wastewater systems is increasing in light of evolving regulatory frameworks. Although routine ARG removal is not yet mandated across the European Union, the 2024 revision of the Urban Waste Water Directive (UWWD)^63^ requires implementation of AMR monitoring systems in large settlements by 2026. Such regulatory developments may ultimately lead to defined resistance-related discharge thresholds, necessitating improved post-treatment solutions. In this context, technologies that achieve substantial microbial and resistome reduction without promoting pathogen enrichment, co-selection, or enhanced mobility represent strategically valuable additions to wastewater treatment infrastructure.

Taken together, this study demonstrates that spatial confinement of disinfectant bioavailability can decouple antimicrobial efficacy from classical resistance selection pathways. By combining robust microbial inactivation, significant ARG and integron reduction, suppression of horizontal gene transfer, and functional reusability without detectable chemical release, immobilised BDMDAC particles could provide a mechanistically distinct framework for AMR-conscious wastewater polishing strategies in the future.

## 5. Acknowledgements

A.E. & T.U.B were supported through the PRESAGE project funded by the Bundesministerium für Forschung, Technologie und Raumfahrt (Grant number 02WAP1619) and by the RHUMARGE project funded by the Deutsche Forschungsgemenischaft (Grant number 544004729). U.K. & T.U.B were supported by the Explore-AMR project and the JPIAMR SEARCHER project funded by the German Bundesministerium für Forschung, Technologie und Raumfahrt under grant numbers 01DO2200 & 01KI24O4A. M.R. was supported through FCT Aquatic/0007/2020 and ID 375 of JPIERANET Aquatic Pollutants, co-financed by ERA-NET Cofund Aquatic Pollutants national funds through FCT/MECI: LEPABE, UID/00511/2025 (https://doi.org/10.54499/UID/00511/2025) and UID/PRR/00511/2025 (https://doi.org/10.54499/UID/PRR/00511/2025) and ALiCE, LA/P/0045/2020 (https://doi.org/10.54499/LA/P/0045/2020). M.R. would like to thank the Foundation for Science and Technology (FCT) for the financial support of the PhD grant (2024.04956.BD). Responsibility for the information and views expressed in the manuscript lies entirely with the authors.

## 6. Author contributions

A.E, M.R., L.M., A.P, U.K. and T.U.B. conceived the study. M.R., A.P and L.M. developed the functionalised particles. M.R. carried out growth assays, plasmid transfer assays and gene quantification through qPCR. A.E. carried out the bioinformatic analysis. M.R., U.K. and A.E. analysed and interpreted the data. M.R and A.E. wrote the initial draft of the manuscript. All authors edited the manuscript and approved its final version.

## 7. Data availability

All raw reads corresponding to 16S rRNA gene sequencing have been deposited and made publicly available at the Short Read Archive (SRA) of the National Center for Biotechnology Information (NCBI) under the BioProject with accession number PRJNA1431717

