## Supplementary_material for "Reusable immobilised quaternary ammonium particles reduce microbial and resistome burdens without promoting resistance selection during wastewater post-treatment"

### Supplementary Information

| Primer name | Sequence (5' - 3') |
|---|---|
| 331F | TCCTACGGGAGGCAGCAGT |
| 518R | ATTACCGCGGCTGCTGG |

| Temperature | Time | |
|---|---|---|
| 95 °C | 5 min | |
| 95 °C | 15 sec | x 45 |
| 60 °C | 1 min | x 45 |
| 72 °C | 30 sec | x 45 |
| 72 °C | 5 min | |

Table S1: Primers and conditions used for 16S rRNA gene quantification

| Reagents | mg/L |
|---|---|
| Peptone | 256.55 |
| Triptone | 354.24 |
| NaCl | 407.4 |
| $Na_2SO_4$ | 44.6 |
| $K_2HPO_4$ | 44.6 |
| $MgCl_2 \bullet 6H_2O$ | 3.7 |
| $FeCl_2 \bullet$2H2O | 3.7 |
| $CaCl_2 \bullet 2H_2O$ | 3.7 |
| $MnSO_4$ | 0.057 |
| $H_2MoO_4$ | 0.031 |
| NaOH | 0.008 |
| $ZnSO_4$ | 0.046 |
| $CoSO_4$ | 0.049 |
| $CuSO_4$ | 0.079 |

Table S2: Synthetic wastewater medium composition

| Target | Forward primer (5'-3') | Reverse primer (5'-3') |
|---|---|---|
| *aac-(3)-VI* | CGTCACTTATTCGATGCCCTTAC | GTCGGGCGCGGCATA |
| *aph-(3')-VIa* | TCTCATGGCGATATCACGGATAG | TTTCCTCCGATGCATCCTCTC |
| *aph-(3'')-Ib* | AACAGGTTTGGGAGGCGATG | CGCAACAAGCCTCTCCTGAA |
| *aac-(6')-Ib* | CGTCGCCGAGCAACTTG | CGGTACCTTGCCTCTCAAACC |
| *aadA1* | GTTGTGCACGACGACATCATT | GGCTCGAAGATACCTGCAAGAA |
| *mcr-1* | CACATCGACGGCGTATTCTG | CAACGAGCATACCGACATCG |
| *bla*$_{\text{CMY\_2}}$ | AAAGCCTCAT GGGTGCATAAA | ATAGCTTTTGTTTGCCAGCATCA |
| *bla*$_{\text{CTX-M}}$ | CGTACCGAGCCGACGTTAA | CAACCCAGGAAGCAGGCA |
| *bla*$_{\text{OXA-48}}$ | TGTTTTTGGTGGCATCGAT | GTAAMRATGCTTGGTTCGC |
| *bla*$_{\text{OXA-58}}$ | GCAATTGCCTTTTAAACCTGA | CTGCCTTTTCAACAAAACCC |
| *bla*$_{\text{TEM}}$ | CGCCGCATACACTATTCTCAG | GCTTCATTCAGCTCCGGTTC |
| *bla*$_{\text{amp-C}}$ | AACAAAAGATCCCCGGTATGG | ACGCCCGTAAATGTTTTGCT |
| *bla*$_{\text{KPC-2}}$ | GCCGCCGTGCAATACAGT | GCCGCCCAACTCCTTCA |
| *dfrA1* | GGAATGGCCCTGATATTCCA | AGTCTTGCGTCCAACCAACAG |
| *ermF* | CAGCTTTGGTTGAACATTTACGAA | AAATTCCTAAAATCACAACCGACAA |
| *ermB* | GAACACTAGGGTTGTTCTTGCA | CTGGAACATCTGTGGTATGGC |
| *mphA* | TCAGCGGGATGATCGACTG | GAGGGCGTAGAGGGCGTA |
| *intI*1 | CGAACGAGTGGCGGAGGGTG | TACCCGAGAGCTTGGCACCCA |
| *mexB* | CTGGAGATCGACGACGAGAAG | GAAATCGTTGACGTAGCTGGAA |
| *qacA* | AAGGGCCACTGCATTAGCTG | CCAGTCCAATCATGCCTGCA |
| *qacH* | CATCGTGCTTGTGGCAGCTA | TGAACGCCCAGAAGTCTAGTTTT |
| *qnrA* | AGGATTTCTCACGCCAGGATT | CCGCTTTCAATGAAACTGCAA |
| *qnrB* | GCGACGTTCAGTGGTTCAGA | GCTGCTCGCCAGTCGAA |
| *qnrS1* | CCACTTTGATGTCGCAGATCTTC | CCCTCTCCATATTGGCATAGGAAA |
| *sul1* | GCCGATGAGATCAGACGTATTG | CGCATAGCGCTGGGTTTC |
| *tetM* | GGAGCGATTACAGAATTAGGAAGC | TCCATATGTCCTGGCGTGTC |
| *tetW* | ATGAACATTCCCACCGTTATCTTT | ATATCGGCGGAGAGCTTATCC |
| *vanA* | GGGCTGTGAGGTCGGTTG | TTCAGTACAATGCGGCCGTTA |
| *incP* | CAGCCTCGCAGAGCAGGAT | CAGCCGGGCAGGATAGGTGAAGT |
| *incW* | AGCGTATGAAGCCCGTGAAGGG | AAAGATAAGCGGCAGGACAATAACG |
| *inuB* | GGATCGTTTACCAAAGGAGAAGG | AGCATAGCCTTCGTATCAGGAA |
| *tolC* | CAGGCAGAGAACCTGATGCA | CGCAATTCCGGGTTGCT |

Table S3: List of selected primers including major ARGs, QAC resistance genes, widespread plasmid markers and genes of taxonomic value for the identification of specific pathogens.

| | Treatment | *k* | *r* | Logistic AUC | Standard error |
|---|---|---|---|---|---|
| Control *E. coli* | 0 mg/mL | 3.288 | 0.011 | 159.788 | 0.164 |
| | 25 mg/mL | 1.586 | 0.01 | 735.133 | 0.037 |
| | 100 mg/mL | 0.304 | 0.007 | 125.789 | 0.023 |
| *E. coli* pKJK5 | 0 mg/mL | 3.053 | 0.012 | 1,507.423 | 0.144 |
| | 25 mg/mL | 1.605 | 0.01 | 741.974 | 0.047 |
| | 100 mg/mL | 0.404 | 0.005 | 93.897 | 0.021 |
| *E. coli* pRP4 | 0 mg/mL | 2.963 | 0.014 | 1,447.326 | 0.111 |
| | 25 mg/mL | 1.42 | 0.01 | 649.155 | 0.027 |
| | 100 mg/mL | - | - | - | - |
| *Control P. putida* | 0 mg/mL | 2.843 | 0.017 | 1,178.49 | 0.087 |
| | 50 mg/mL | 0.719 | 0.028 | 309.396 | 0.045 |
| | 200 mg/mL | - | - | - | - |
| *P. putida* pKJK5 | 0 mg/mL | 3.173 | 0.015 | 1,222.449 | 0.066 |
| | 50 mg/mL | 0.451 | 0.046 | 179.477 | 0.05 |
| | 200 mg/mL | - | - | - | - |
| *P. putida* pRP4 | 0 mg/mL | 3.241 | 0.009 | 988.364 | 0.122 |
| | 50 mg/mL | 0.558 | 0.028 | 160.549 | 0.026 |
| | 200 mg/mL | - | - | - | - |

Table S4: Growth curve parameters for *E. coli* and *P. putida* exposed to BDMDAC-FPs. *k*=Theoretical carrying capacity, *r*= Growth rate. Where no values are given, a logistic curve could not be fit.

| | Treatment | *k* | *r* | Logistic AUC | Standard error |
|---|---|---|---|---|---|
| Control *E. coli* | 0 mg/mL | 3.100 | 0.013 | 1,479.057 | 0.123 |
| | 25 mg/mL | - | - | - | - |
| | 100 mg/mL | - | - | - | - |
| *E. coli* pKJK5 | 0 mg/mL | 3.116 | 0.011 | 1,474.482 | 0.154 |
| | 25 mg/mL | - | - | - | - |
| | 100 mg/mL | - | - | - | - |
| *E. coli* pRP4 | 0 mg/mL | 3.195 | 0.013 | 1,577.850 | 0.137 |
| | 25 mg/mL | - | - | - | - |
| | 100 mg/mL | - | - | - | - |
| Control *P. putida* | 0 mg/mL | 3.054 | 0.017 | 1,488.668 | 0.090 |
| | 50 mg/mL | - | - | - | - |
| | 200 mg/mL | - | - | - | - |
| *P. putida* pKJK5 | 0 mg/mL | 3.158 | 0.015 | 1,452.215 | 0.091 |
| | 50 mg/mL | - | - | - | - |
| | 200 mg/mL | - | - | - | - |
| *P. putida* pRP4 | 0 mg/mL | 2.946 | 0.017 | 1,427.481 | 0.107 |
| | 50 mg/mL | - | - | - | - |
| | 200 mg/mL | - | - | - | - |

Table S5: Growth curve parameters for *E. coli* and *P. putida* exposed to reused BDMDAC-FPs (single reuse). *k*=Theoretical carrying capacity, *r*= Growth rate. Where no values are given, a logistic curve could not be fit.
